# What makes an angry face uncomfortable? Distinct contributions of facial expression, interpersonal distance, facial stimulus type, and gaze in virtual reality

**DOI:** 10.64898/2026.08.06.743152

**Authors:** Hatem Dahech, Tetsuto Minami, Shigeki Nakauchi, Hideki Tamura

## Abstract

Why does an angry face feel uncomfortable? The answer is that it signals a threat. However, a face is only part of an encounter, and distance, facial stimulus type, and gaze may shape discomfort regardless of perceived anger. To separate these cues, we conducted three within-subjects virtual reality experiments. In each experiment, 24 adults viewed avatars at intimate, personal, and social distances (30, 100, and 300 cm, respectively) and rated the face’s perceived anger and their own discomfort; head movement was recorded in Experiments 2 and 3. In Experiment 1, the expression (angry, neutral) and facial color (natural, red) were crossed with distance; in Experiment 2, a featureless mannequin served as a nonface comparison; and in Experiment 3, the gaze direction (direct, averted) was manipulated. Expression primarily determined perceived anger, whereas distance predominantly determined discomfort: A nearby neutral face was uncomfortable despite low perceived anger (Experiment 1). A neutral human face was more uncomfortable than a mannequin, although both received similarly low perceived-anger ratings (Experiment 2). Direct gaze increased the discomfort without changing perceived anger (Experiment 3). Backward head movement exhibited a similar pattern, with participants leaning back more from human faces than from the mannequin. These results indicate that the discomfort associated with an angry face is not merely explained by perceived anger. Instead, social discomfort was differentially associated with interpersonal distance and gaze direction and differed between the human-face and mannequin conditions.

## 1. Introduction

Among facial expressions, anger in particular signals a possible threat. Angry faces are detected rapidly and more efficiently than neutral or happy faces are (Fox et al., 2000; Öhman et al., 2001), and electrophysiological studies have reported earlier and stronger attentional allocation to angry faces than to happy faces (Feldmann-Wüstefeld et al., 2011). Beyond the context of detection, angry faces shape how people act toward one another, as people initiate movement away from angry faces more readily than toward them (Stins et al., 2011; Paulus & Wentura, 2016), and they maintain a greater interpersonal distance from angry others than from friendly or neutral others, both in person and in virtual reality (Bönsch et al., 2018; Ruggiero et al., 2017; Meinhardt-Injac et al., 2025). Recent VR studies have demonstrated that angry expressions can interfere with an ongoing perceptual task and that spatial distance and emotional expression jointly shape visual attention toward different regions of a virtual human’s body (Rapuano et al., 2023, 2026). How angry a face appears can also be influenced by lower-level cues: a reddish facial color biases observers toward perceiving anger and increases the rated strength of the expression (Nakajima et al., 2017; Minami et al., 2018; Peromaa & Olkkonen, 2019; Shibusawa et al., 2025). Related research has demonstrated that facial color influences implicit facial expression processing, facial expression influences facial color memory, and facial expression and color interact in the P3 component of the event-related potential component (Nguyen et al., 2023; Hasegawa et al., 2024, 2025). Because angry faces signal threats and produce avoidance, it is typically assumed that the discomfort associated with encountering an angry face is a direct consequence of perceiving its threat. In other words, the face is taken to be uncomfortable because it is angry.

However, a social encounter involves more than merely a face. Interpersonal distance itself is a powerful determinant of how an encounter feels. Hall (1966) described how people divide the space surrounding the body into intimate, personal, and social zones, and Argyle and Dean (1965) proposed that social closeness is actively regulated through distance, eye contact, and other cues. Studies that have relied on virtual humans have further demonstrated that interpersonal comfort distance and reaching distance vary with different tasks and interaction conditions and revealed that spatial behavior is shaped by social information such as moral impressions, gender, and age (Iachini et al., 2014, 2015, 2016). When another person enters their personal space, people report experiencing discomfort and physiological arousal even in the absence of any threatening expression (Hayduk, 1983; Candini et al., 2021). Angry virtual humans have been reported to increase both interpersonal comfort distance and reaching distance and to elicit stronger psychophysiological responses than neutral or happy virtual humans (Ruggiero et al., 2021). The amygdala is more active when another person is close than when that person is far, and a patient with bilateral amygdala damage reported no discomfort even at a nose-to-nose distance (Kennedy et al., 2009). The regulation of interpersonal distance is a distinct process that varies across individuals and is altered in clinical conditions (Welsch et al., 2020). These findings indicate that proximity alone can elicit discomfort, thus raising the possibility that the discomfort of an angry face does not depend solely on its expression. A neutral face may feel uncomfortable when it is very close, whereas an angry face may feel only mildly uncomfortable when it is far away.

If proximity can produce discomfort on its own, other social cues may do so as well. One such cue is the social nature of the stimulus. A close human face may be uncomfortable not only because it is near but also because its human facial appearance may carry social meaning that is absent from a nonhuman object at the same distance (Bailenson et al., 2001). A second cue is gaze direction. Direct gaze indicates that the other person’s attention is directed at the observer and thus increases the self-relevance of the encounter, and direct and averted gaze engage the approach and avoidance systems differently (Hietanen et al., 2008). Gaze has also been reported to interact with expression, as some studies that have reported that gaze can shift the perceived emotion of a face, particularly in cases of anger (Adams & Kleck, 2005; Ewbank et al., 2009; McCrackin & Itier, 2019). Critically, these cues may change how uncomfortable an encounter feels without changing how angry the face is perceived to be. A neutral face is no angrier as a result of being a human face rather than a nonhuman object, and an angry face is no angrier if it looks at the observer rather than away; however, either case may feel more uncomfortable.

Taken together, the results of previous studies have suggested that angry expressions, interpersonal distance, and gaze influence social behavior. However, the question of whether the discomfort associated with an angry face is merely a consequence of perceiving anger or whether it is shaped independently by these other social cues remains unanswered. Few studies have measured perceived anger and subjective discomfort within the same paradigm; thus, these two aspects have rarely been separated. Consequently, it remains unknown whether a cue that raises discomfort does so by making the face appear angrier or by acting on the feeling of the encounter directly.

The goal of the present study was to identify the factors that make an angry face uncomfortable. We distinguished between two judgments that are usually combined: the perceived anger of the face, which involves an evaluation of the face itself, and the discomfort involved in the encounter, which represents an evaluation of how the situation felt. We used virtual reality (VR), which allowed us to present the same face at controlled distances and with different gaze directions while holding the rest of the situation constant. VR facilitates a controlled investigation of responses to virtual humans at interpersonal distances (Bailenson et al., 2003; Tootell et al., 2021). In the context of three within-subjects experiments, 24 adults viewed photorealistic avatars at three distances that corresponded to Hall’s intimate, personal, and social zones, and in every trial, they rated the perceived anger of the face on a scale ranging from 1 (not angry) to 10 (very angry); they also rated their own discomfort on a scale ranging from 1 (comfortable) to 10 (uncomfortable). In Experiments 2 and 3, head movement was also recorded in six degrees of freedom as an additional bodily index of the response. The three experiments isolated different cues. Experiment 1 manipulated facial expression (angry, neutral) and color (natural, red) across distances, compared the contribution of facial expression with that of proximity, and tested whether the two judgments followed the same cue. Experiment 2 replaced color with a featureless mannequin that was presented alongside angry and neutral faces, thereby testing whether discomfort differed between human facial stimuli and a head-shaped stimulus that lacked facial features at the same distance. Experiment 3 manipulated gaze direction (Direct, Averted), thus isolating the effect of being looked at and testing whether it can elicit discomfort without changing perceived anger.

Our central hypothesis distinguished between two accounts. If discomfort is solely a consequence of perceived anger, then any manipulation that increases discomfort should also increase perceived anger, and these two factors should rise and fall together. Alternatively, if discomfort is shaped by social cues that extend beyond facial expression, interpersonal distance, facial stimulus type, and gaze should influence discomfort without necessarily increasing perceived anger.

### 2. Experiment 1

Experiment 1 examined whether social discomfort can be explained solely by perceived anger or whether interpersonal distance makes an independent contribution in this context. If discomfort is merely a consequence of perceiving anger, it should primarily follow facial expression. Alternatively, if interpersonal distance independently shapes discomfort, close interpersonal distance should increase discomfort even when the level of perceived anger is low. Previous studies have reported that angry faces are detected rapidly and efficiently (Fox et al., 2000; Öhman et al., 2001), whereas personal-space intrusion is associated with discomfort and physiological arousal (Hayduk, 1983; Kennedy et al., 2009; Candini et al., 2021). Therefore, we hypothesized that perceived anger is driven primarily by facial expression, whereas discomfort is driven primarily by interpersonal distance. Specifically, a neutral face was expected to elicit greater discomfort at close distances, whereas an angry face was expected to elicit relatively low levels of discomfort at far distances. In addition, we manipulated facial color (natural vs. red) because facial redness has been reported to enhance perceptions of anger (Nakajima et al., 2017; Minami et al., 2018; Peromaa & Olkkonen, 2019; Shibusawa et al., 2025). This manipulation allowed us to test whether a facial cue that increases perceived anger also increases discomfort and therefore whether these two judgments respond similarly to the same visual cue.

### 2.1. Methods

#### 2.1.1. Participants

Twenty-four adult participants (20 males and 4 females; all Asian) were included in this experiment (mean age = 24 years; *SD* = 1.6). The sample size was determined on the basis of a previous study that examined emotional facial expression and interpersonal comfort distance, which focused on 24 participants (Ruggiero et al., 2021). An a priori power analysis that was conducted using PANGEA (Power Analysis for General Anova designs (Westfall, 2016)) indicated that 17 participants were required to detect the main effect of distance at a power of.80, assuming an effect size of d = 0.5, α =.05, and four replicates per condition. All the participants provided written informed consent prior to their participation in the experiment. This study was approved by the Institutional Review Board of Toyohashi University of Technology (approved numbers 2024-28 and 2025-31).

#### 2.1.2. Apparatus

The experimental stimuli were presented using an HTC VIVE Pro Eye head-mounted display (HMD; resolution = 1440 × 1600 pixels per eye, refresh rate = 90 Hz, field of view = 110°), supported by a Windows PC. The experiment was developed in Unreal Engine 5.5 and presented through SteamVR. Responses were recorded using a handheld HTC VIVE controller.

#### 2.1.3. Stimuli

The virtual humans used in the experiment were photorealistic, full-bodied avatars that were created using the web-based MetaHuman Creator associated with Unreal Engine 5.5; each avatar was rendered as a standing figure with natural skin detail (Figure 1). Four East Asian avatar identities were used, including two males and two females. The avatars were designed to appear as adults; no numerical age was assigned, and age was not manipulated experimentally. To avoid differences unrelated to facial identity, all the avatars were presented with the same hairstyle, clothing, and body shape.

**Figure 1.**
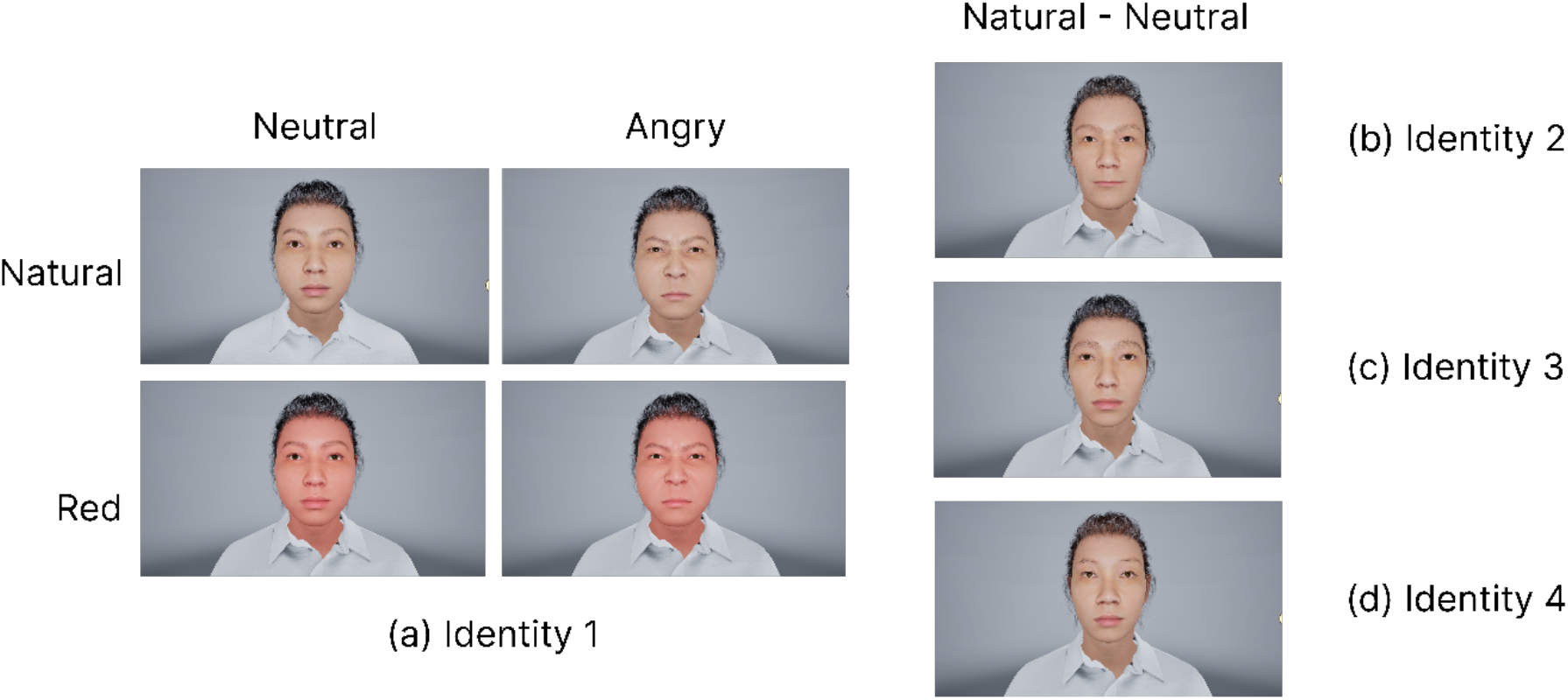
Stimuli used in Experiment 1. Each identity was presented with angry and neutral expressions in both natural and red color at three distances: 30, 100, and 300 cm.

Each avatar featured one of two facial expressions: angry or neutral. Within each identity, the same MetaHuman asset was used for both expressions as follows. The angry expression was created using the MetaHuman Facial Pose Library preset “Anger-03_Anger”, whereas the neutral expression used the character’s default facial pose; no expression preset was applied. Thus, expression was manipulated by changing only the facial expression preset; identity, hairstyle, clothing, body shape, and skin texture were held constant. Expressions were based on MetaHuman facial expression presets that were applied to the facial rig. The angry preset mainly involved lowered and tightened brows, narrowed eyes, and increased tension both around the mouth and above the nose. Additionally, because the mouth shape used in the original angry preset produced an unnatural open mouth shape, it was manually adjusted. Other than the mouth shape of the angry preset, no other facial features were manually modified. The neutral preset kept the face relaxed without any clear emotional expression.

Facial color was also manipulated between a natural skin tone (natural) and a red tint (red). The red tint was created by increasing the redness of the skin by approximately 12 units (Nakajima et al. 2017) on the a* (red-green) axis of the CIELAB color space (mean Δ*a*^∗^ = 12.30, range 12.04–12.64).

#### 2.1.4. Procedure

The procedure used for Experiment 1 is illustrated in Figure 2. The participants were seated and wore an HMD with hand-held controllers (Figure 2A). Each trial (Figure 2B) proceeded as follows: the participant initiated the trial when ready by clicking the “Start Experiment Button”; a single avatar then appeared at one of the three distances (30, 100, and 300 cm) that corresponded to Hall’s intimate, personal, and social zones (Hall 1966) and remained visible for 10 s at eye level at one of three distances from the participant. Subsequently, a rating screen was presented, on which the participants rated the perceived anger of the facial expressions (1 = not angry, 10 = very angry) and their own discomfort (1 = comfortable, 10 = uncomfortable). The design crossed expression (Angry, Neutral) × color (Natural, Red) × distance (30, 100, and 300 cm), resulting in 12 conditions. Each condition was repeated across 4 identities, for a total of 48 trials per participant, which were presented in random order. No scheduled rest breaks were included. However, participants initiated each trial themselves and could pause before starting the next trial if needed.

**Figure 2.**
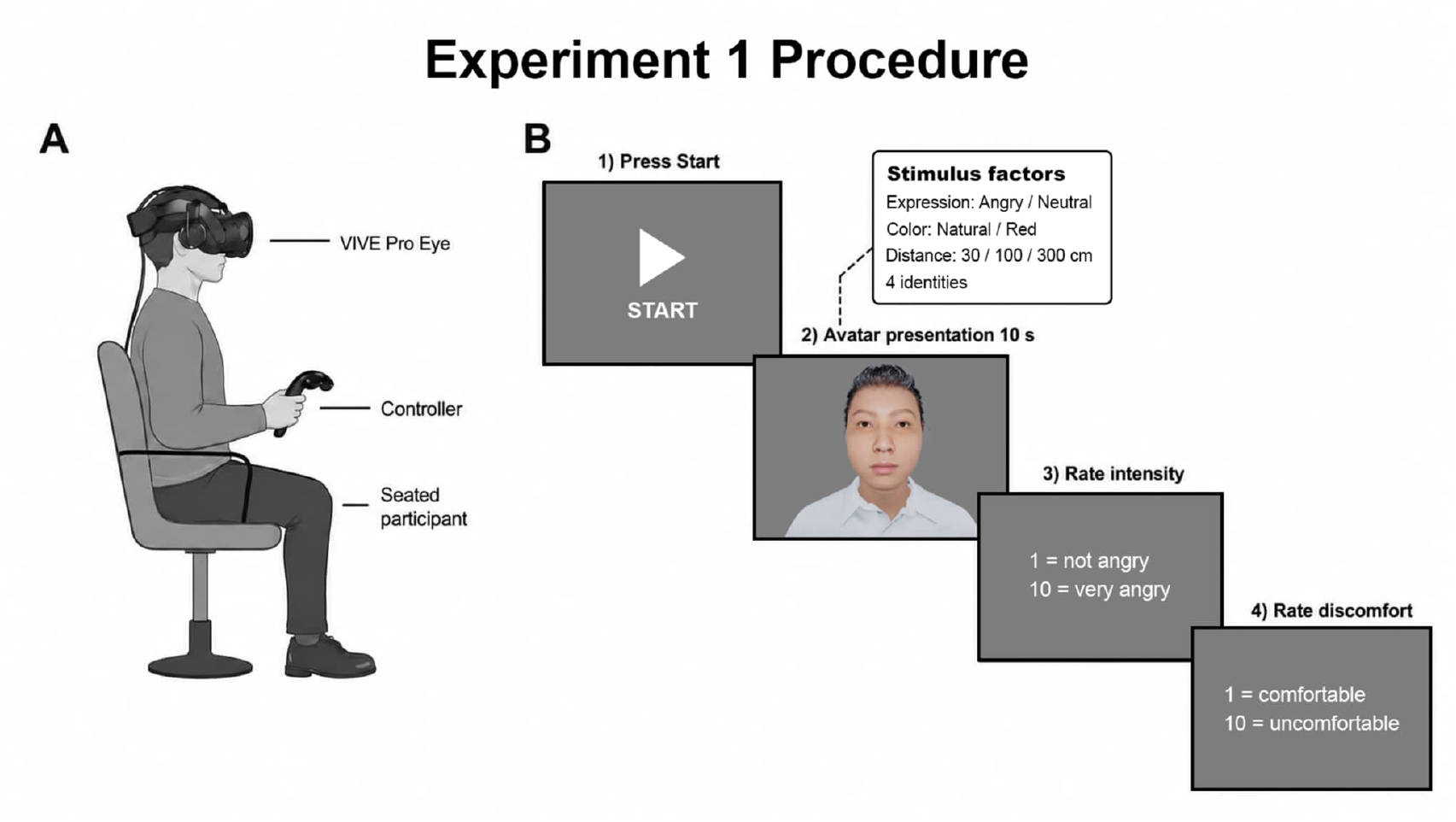
Experiment 1: Trial procedure. (A) Experimental setup. (B) Trial procedure. The participant initiated the trial, the avatar appeared at one of the three distances for 10 s, and the participant then rated perceived anger and discomfort.

#### 2.1.5. Data analysis

Statistical analyses were performed in Python using the “statsmodels” and “scipy” packages. The perceived anger and discomfort ratings were analyzed by conducting a three-way repeated-measures analysis of variance (ANOVA), in which expression, color, and distance served as within-participant factors. For factors that featured more than two levels, Greenhouse–Geisser-corrected degrees of freedom and p values were reported when the assumption of sphericity was violated. Effect sizes are reported as partial eta-squared (*η_p_*^2^). Significant effects were followed up by performing pairwise *t* tests, and the Bonferroni correction was applied for multiple comparisons.

### 2.2. Results

#### 2.2.1. Perceived anger

The results of the perceived-anger ratings are presented in Figure 3. We performed a three-way repeated-measures ANOVA on the perceived-anger ratings (Table S1). A significant main effect of expression was observed, *F*(1, 23) = 582.07, *p* <.001, *η_p_*^2^ =.962; angry faces were rated as much angrier than neutral faces were, *t*(23) = 24.13, *adj*. *p* <.001, *d* = 4.92. The main effect of distance was also significant, *F*(1.21,27.86) = 73.81, *p* <.001, *η_p_*^2^ =.762; all three distances differed, and closer faces were rated as angrier (30 vs. 100 cm, *t*(23) = 8.75, *adj*. *p* <.001 *d* = 1.78; 30 vs. 300 cm, *t*(23) = 9.02, *adj*. *p* <.001, *d* = 1.84; 100 vs. 300 cm, *t*(23) = 6.96, *adj*. *p* <.001, *d* = 1.42). Color had a significant main effect, *F*(1,23) = 29.95, *p* <.001, *η_p_*^2^ =.566; red faces were rated as angrier than natural faces were, *t*(23) = 5.47, *adj*. *p* <.001, *d* = 1.12).

**Figure 1.**
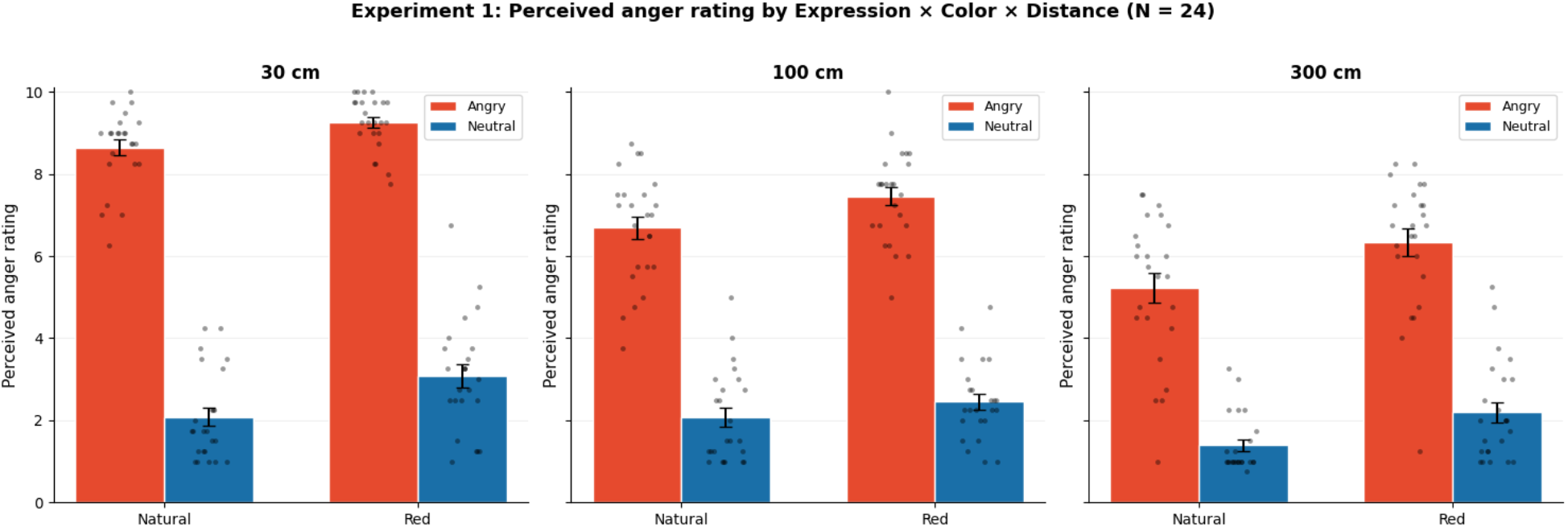
Perceived-anger ratings in Experiment 1 as a function of facial expression (angry, neutral), facial color (natural, red), and distance (30, 100, and 300 cm). The bars represent condition means, the error bars represent standard errors, and the points represent individual participant values.

Additionally, the distance × expression interaction was significant, *F*(2, 46) = 49.37, *p* <.001, *η_p_*^2^ =.682: the difference between angry and neutral faces was greatest at the closest distance and decreased with increasing distance, although it remained large throughout (30 cm, *t*(23) = 30.21, *d* = 6.17; 100 cm, *t*(23) = 20.00, *d* = 4.08; 300 cm, *t*(23) = 13.40, *d* = 2.73; *and all adj*. *p* <.001. In contrast, the distance × color interaction, *F*(2, 46) = 1.89, *p* =.163, *η_p_*^2^ =.076, and the expression × color interaction, *F*(1,23) = 0.37, *p* =.547, *η_p_*^2^ =.016, were not significant. The three-way interaction was significant, *F*(2,46) = 3.27; *p* =.047, *η_p_*^2^ =.125: red increased perceived anger in every expression-by-distance cell with the exception of neutral faces at 100 cm (*t*(23) = 1.64, *adj*. *p* =.688 *d* = 0.33), and its effect was strongest for angry faces at the farthest distance (*t*(23) = 6.17, *adj*. *p* <.001 *d* = 1.26) (Table S2). Among these effects, expression was by far the strongest driver of perceived anger.

#### 2.2.2. Discomfort

The results of the perceived discomfort ratings are presented in Figure 4. The same three-way ANOVA was performed on the discomfort ratings (Table S3). The main effect of distance was significant, *F*(2,46) = 236.47, *p* <.001, *η_p_*^2^ =.911, as closer faces were rated as more uncomfortable (30 vs. 100 cm, *t*(23) = 15.24, *adj*. *p* <.001 *d* = 3.11; 30 vs. 300 cm, *t*(23) = 18.24, *adj*. *p* <.001 *d* = 3.72; 100 vs. 300 cm, *t*(23) = 8.63, *adj*. *p* <.001 *d* = 1.76). The main effect of expression was also significant: *F*(1,23) = 79.96, *p* <.001, *η_p_*^2^ =.777. Angry faces were rated as more uncomfortable than neutral faces were, *t*(23) = 8.94, *adj*. *p* <.001, *d* = 1.83. Color also had a significant main effect, *F*(1,23) = 18.05, *p* <.001, *η_p_*^2^ =.440. Red faces were rated as more uncomfortable than natural faces were. *t*(23) = 4.25, *adj*. *p* <.001 *d* = 0.87.

**Figure 4.**
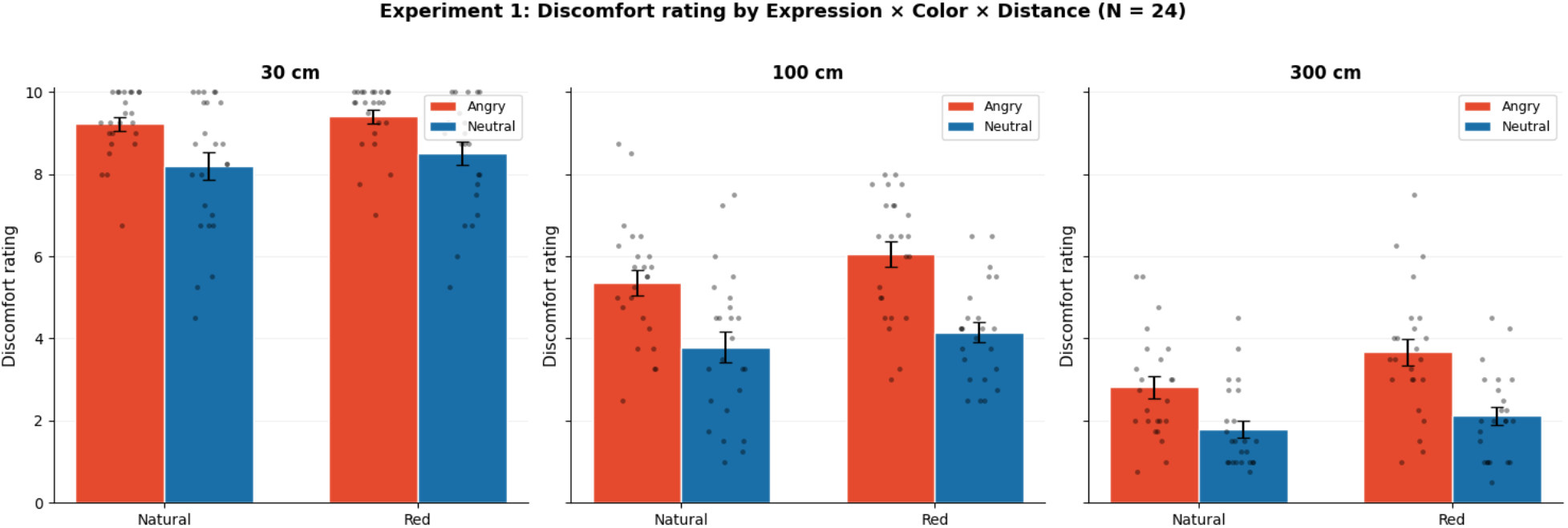
Discomfort ratings in Experiment 1 as a function of facial expression (Angry, Neutral), facial color (natural, red), and distance (30, 100, and 300 cm). The bars represent condition means, the error bars represent standard errors, and the points represent individual participant values.

Additionally, the distance × expression interaction was significant, *F*(2, 46) = 4.25, *p* =.020, *η_p_*^2^ =.156: unlike in the case of perceived anger, the difference between angry and neutral faces increased with distance (30 cm, *t*(23) = 4.68, *adj*. *p* <.001 *d* = 0.95; 100 cm, *t*(23) = 6.78, *adj*. *p* <.001 *d* = 1.38; 300 cm, *t*(23) = 7.49, *adj*. *p* <.001 *d* = 1.53). The expression × color interaction was also significant, *F*(1,23) = 4.33, *p* =.049, *η_p_*^2^ =.159: red amplified the discomfort associated with angry faces (*t*(23) = 4.90, *adj*. *p* <.001, *d* = 1.00) to a greater extent than that associated with neutral faces (*t*(23) = 2.66, *adj*. *p* =.028, *d* = 0.54). In contrast, the distance × color interaction, *F*(2,46) = 2.75, *p* =.075, and *η_p_*^2^ =.107, and the three-way interaction, *F*(2,46) = 3.08, *p* =.055, and *η_p_*^2^ =.118, were not significant. Unlike in the case of perceived anger, the strongest driver of discomfort was distance rather than expression.

### 2.3. Discussion

In Experiment 1, participants rated the perceived anger and discomfort associated with angry and neutral faces, which were shown in a natural or red color, at three distances. As indicated by the corresponding ANOVAs, expression had the strongest effect on perceived-anger ratings, whereas distance had the strongest effect on discomfort ratings. This pattern suggests that perceived anger and discomfort judgments were differentially sensitive to expression and distance: how angry a face appears depends primarily on its expression, whereas how uncomfortable the encounter feels depends primarily on proximity. These results are consistent with previous findings indicating that angry faces are detected rapidly as threat-relevant signals (Fox et al., 2000; Öhman et al., 2001) and that close interpersonal distance produces discomfort and physiological arousal (Hayduk, 1983; Kennedy et al., 2009; Candini et al., 2021). Additionally, the neutral face at the closest distance illustrates this point directly: it was rated low in perceived anger but high in discomfort. Therefore, we suggest that a face can be felt as uncomfortable without being perceived as angry, which is consistent with the claim that proximity contributes to discomfort independently of expression (Hayduk, 1983; Kennedy et al., 2009; Candini et al., 2021).

However, in Experiment 1, a human face was present in every condition. The question whether the discomfort observed with regard to human faces also occurs when a head-shaped stimulus that lacks human facial features remains unanswered. To address this question, in Experiment 2, we used a featureless mannequin alongside the angry and neutral faces.

### 3. Experiment 2

Experiment 2 examined whether discomfort differed between a human facial stimulus and a featureless mannequin that were presented at the same distance. Because all the stimuli used in Experiment 1 were human faces, the question of whether the observed discomfort would also have occurred for a head-shaped stimulus that lacked facial features remained unanswered. We therefore introduced a featureless mannequin alongside the angry and neutral faces. We predicted that a neutral human face would elicit greater discomfort than the mannequin would despite similarly low perceived-anger ratings. This comparison tested whether discomfort differed as a function of stimulus type, but it was not intended to serve as a direct manipulation of social or human presence.

We also recorded head movements because avoidance responses may be expressed not only through explicit judgments but also through subtle bodily movements, such as leaning away from a stimulus (Stins et al., 2011; Welsch et al., 2023). We therefore expected to observe backward head displacement, which was used as an additional behavioral measure, to broadly parallel subjective discomfort.

### 3.1. Methods

#### 3.1.1. Participants

Twenty-four participants participated in this experiment (21 males and 3 females; all Asian). Mean age = 22.3 years, *SD* = 1.3. Among these individuals, six participants had also completed Experiment 1. All other participant details, including the consent procedure and ethics approval, were the same as those described for Experiment 1.

#### 3.1.2. Apparatus

The apparatus used in this experiment was identical to that employed for Experiment 1. Head position and orientation, as recorded by the HMD, were logged in six degrees of freedom throughout each trial.

#### 3.1.3. Stimuli

Three face types were used: angry, neutral, and a mannequin. The angry and neutral avatars were the same MetaHuman faces that were used in Experiment 1. The mannequin was a featureless head with no facial features (Figure 5), which was matched to the avatars in terms of both size and position. As in Experiment 1, each stimulus was presented at eye level at one of three distances from the participant (30, 100, or 300 cm).

**Figure 5.**
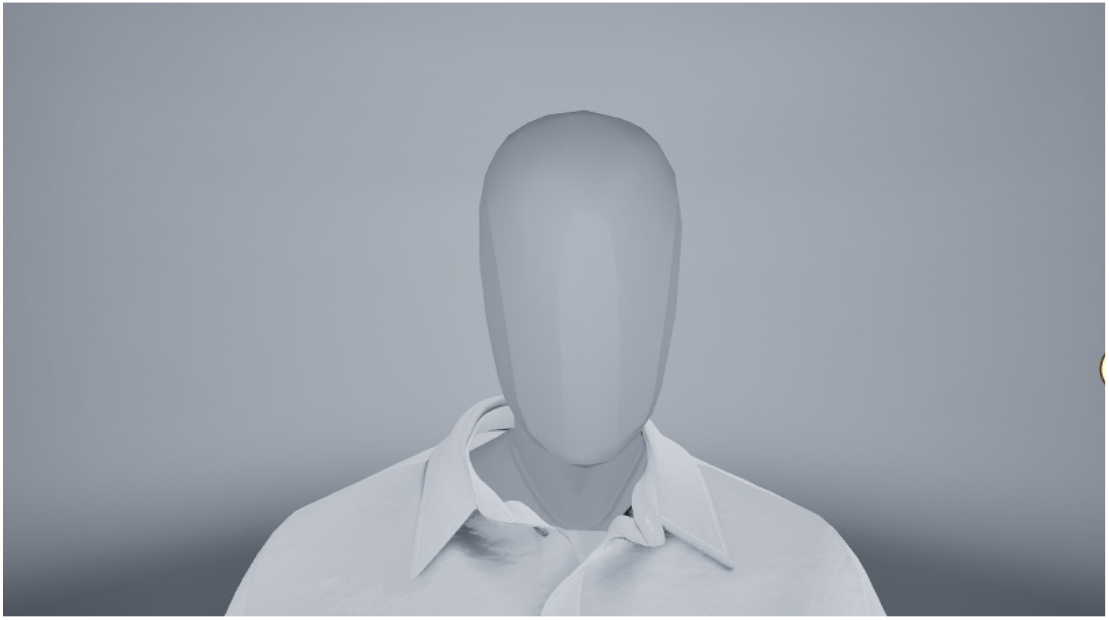
Mannequin stimulus used in Experiment 2.

#### 3.1.4. Procedure

The procedure used in this experiment was identical to that employed for Experiment 1. The design crossed face type (angry, neutral, mannequin) × distance (30, 100, 300 cm), resulting in 9 conditions. For the angry and neutral face conditions, each distance was presented with four avatar identities and repeated twice. With respect to the mannequin condition, the same mannequin was presented at each distance and repeated the same number of times as each human face type. This approach resulted in 24 trials per face type and 72 trials per participant. In each trial, the stimulus was shown for 10 s, after which the participants rated the perceived anger of the face and their own level of discomfort, each of which was scored on a scale ranging from 1 to 10, as in Experiment 1.

#### 3.1.5. Data analysis

In addition to the ratings, head movement was analyzed in this experiment. The head pose was logged in six degrees of freedom, but the present analysis focused on translation along the forward-backward (X) and lateral (Y) axes. For each trial, both axes were baseline corrected by subtracting the value of the first sample, such that all trajectories began at zero. A single peak value was then extracted for each axis. For the X axis, the peak was taken as the most negative baseline value, thus representing backward movement. For the Y axis, the peak was taken as the maximum absolute deviation, thus representing lateral movement. The trials were averaged to one value per participant per cell before analysis. The associations between backward displacement and the two ratings were assessed by reference to repeated-measures correlations, in which context participants were treated as the clustering variable.

### 3.2. Results

#### 3.2.1. Perceived anger

We performed a two-way repeated-measures ANOVA on the perceived-anger ratings (Figure 6; Table S4). A significant main effect of facial type was observed, *F*(1.52,34.86) = 144.04, *p* <.001, *η_p_*^2^ =.862. Post hoc tests revealed that angry faces received considerably higher perceived-anger ratings than both neutral faces, *t*(23) = 14.24, *adj*. *p* <.001, *d* = 2.91, and the mannequin, *t*(23) = 12.45, *adj*. *p* <.001, *d* = 2.54, whereas neutral faces and the mannequin did not differ in this regard, *t*(23) = 1.61, *adj*. *p* =.366, *d* = 0.33. The main effect of distance was also significant, *F*(1.16,26.59) = 64.90, *p* <.001, *η_p_*^2^ =.738, as closer faces were rated as angrier. The face type × distance interaction was significant: *F*(1.88,43.28) = 27.26, *p* <.001, *η_p_*^2^ =.542. Follow-up t tests revealed that perceived-anger ratings were higher for angry faces than for both neutral faces and the mannequin at all distances (*all adj*. *p* <.001), whereas neutral faces and the mannequin did not differ at any distance (*all adj*. *p* ≥.05). These findings indicate that high perceived-anger ratings were specific to angry facial expressions.

**Figure 2.**
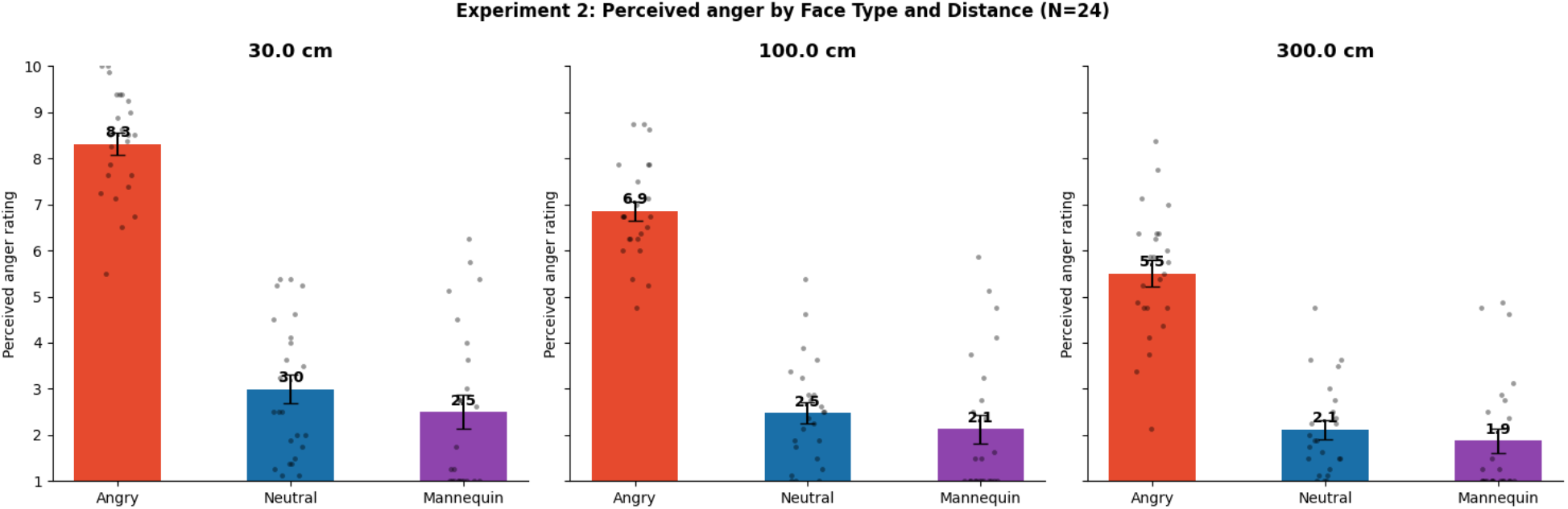
Perceived-anger ratings in Experiment 2 as a function of face type (angry, neutral, mannequin) and distance (30, 100, and 300 cm). The bars represent condition means, the error bars represent standard errors, and the points represent individual participant values.

#### 3.2.2. Discomfort

The same ANOVA was performed on the discomfort ratings (Figure 7; Table S5). The main effect of distance was significant, *F*(1.51,34.77) = 147.59, *p* <.001, *η_p_*^2^ =.865. Pairwise comparisons revealed that all three distances differed, as closer encounters were rated as more uncomfortable: 30 vs. 100 cm, *t*(23) = 10.73, *d* = 2.19; 30 vs. 300 cm, *t*(23) = 14.31, *d* = 2.92; 100 vs. 300 cm, *t*(23) = 7.58, *d* = 1.55; all *adj*. *p* ≤.001. The main effect of face type was also significant, *F*(1.47,33.73) = 75.11, *p* <.001, *η_p_*^2^ =.766. Post hoc tests revealed that all three face types differed. Angry faces were rated as more uncomfortable than neutral faces were, *t*(23) = 9.42, *adj*. *p* <.001, *d* = 1.92. Angry faces were also rated as more uncomfortable than the mannequin was, *t*(23) = 9.41, *adj*. *p* <.001, *d* = 1.92. Unlike in the case of perceived anger, neutral faces were rated as more uncomfortable than the mannequin was, *t*(23) = 4.13, *adj*. *p* =.001, *d* = 0.84.

**Figure 7.**
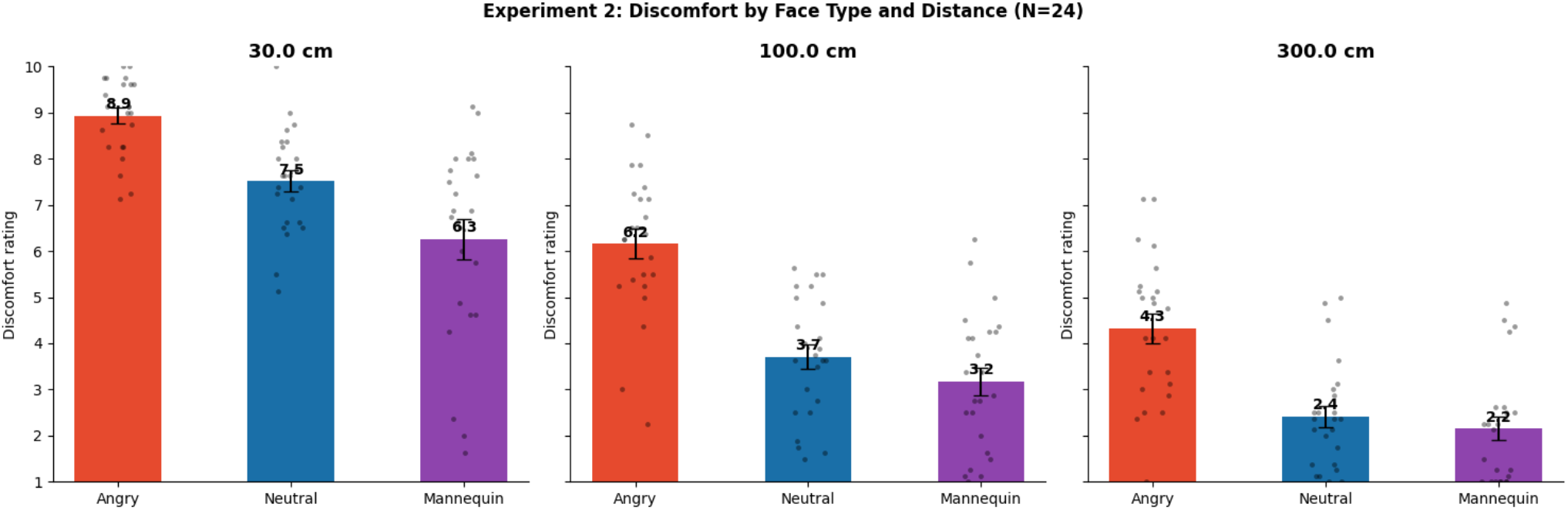
Discomfort ratings in Experiment 2 as a function of face type (angry, neutral, or mannequin) and distance (30, 100, and 300 cm). The bars represent condition means, the error bars represent standard errors, and the points represent individual participant values.

Additionally, the face type × distance interaction was significant, *F*(1.66,38.09) = 5.75, *p* =.010, *η_p_*^2^ =.200. Follow-up t tests revealed that angry faces were rated as more uncomfortable than both neutral faces and the mannequin were at all distances (*all adj*. *p* <.001). Neutral faces were rated as more uncomfortable than the mannequin was at 30 cm, *t*(23) = 4.09, *adj*. *p* =.001, *d* = 0.84, and 100 cm, *t*(23) = 2.88, *adj*. *p* =.025, *d* = 0.59, but not at 300 cm, *t*(23) = 1.36, *adj*. *p* =.564, *d* = 0.28. Thus, a neutral face elicited more discomfort than a featureless mannequin did, mainly at closer distances.

#### 3.2.3. Head movement

For head movement, each trial was baseline corrected to its first sample, and one peak value was extracted for each axis. These peak values were then averaged within each participant and condition and included in the ANOVA (Table S6). In Figure 8, backward movements were significant main effects of face type, *F*(2,46) = 16.51, *p* <.001, *η_p_*^2^ =.418, and distance, *F*(1.39,32.04) = 5.33, *p* =.018, *η_p_*^2^ =.188, and a significant face type × distance interaction, *F*(2.79,64.21) = 4.68, *p* =.006, *η_p_*^2^ =.169. Across distances, both faces produced more backward movement than the mannequin did (Table S7): angry vs. mannequin, *t*(23) = −5.33, *d* = −1.09; mannequin vs. neutral, *t*(23) = 3.73, *d* = 0.76; both *adj*. *p* <.005. Angry and neutral faces did not differ, *t*(23) = −2.29, *adj*. *p* =.095, *d* = −0.47. Because the interaction was significant, we compared face types within each distance. At 30 cm, all three differed: angry faces produced more backward movement than neutral faces did, *t*(23) = −3.23, *adj*. *p* =.011, *d* = −0.66; angry faces produced more such movement than the mannequin did, *t*(23) = −4.76, *adj*. *p* <.001, *d* = −0.97; and neutral faces produced more than the mannequin, did *t*(23) = 3.53, *adj*. *p* =.005, *d* = 0.72. At 100 and 300 cm, the face types were not associated with such differences. Accordingly, backward movement followed the order angry > neutral > mannequin, but only at the closest distance.

**Figure 8.**
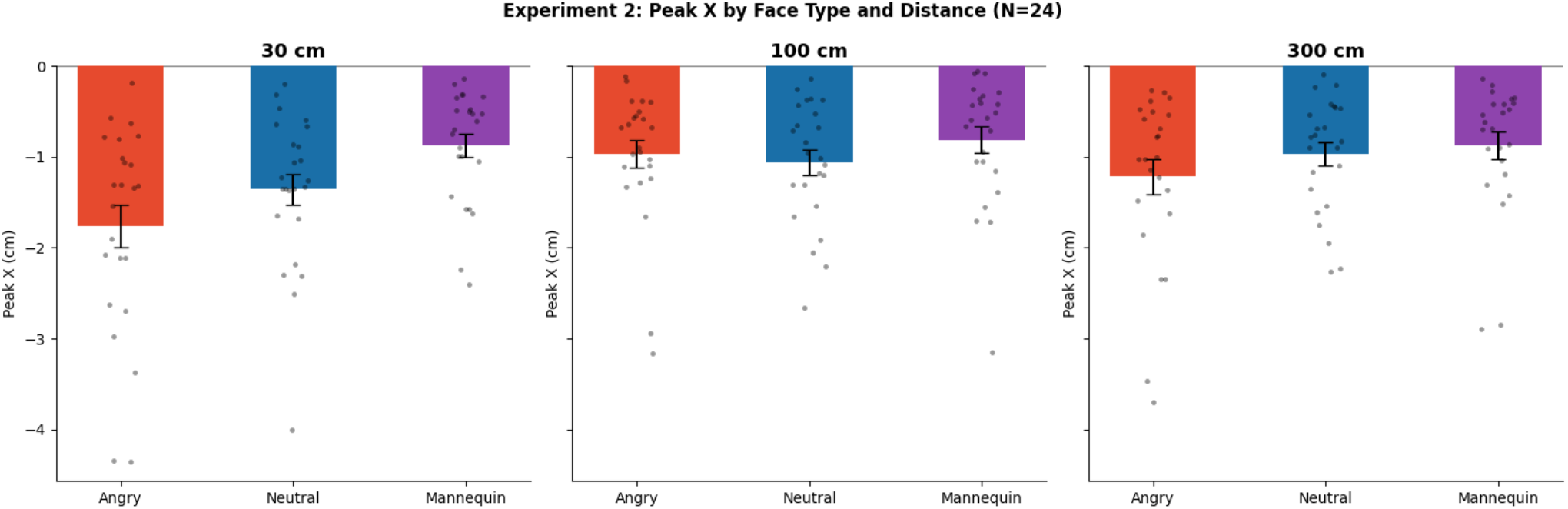
Peak backward head movement in Experiment 2 as a function of face type and distance.

The lateral axis did not demonstrate any significant effects of face type, distance, or their interaction: for lateral movement and face type *F*(2,46) = 1.94, *p* =.155, *η_p_*^2^ =.078, no main effect of distance was observed, *F*(1.33,30.63) = 1.52, *p* =.233, *η_p_*^2^ =.062, nor was that the case for the interaction, *F*(2.85,65.57) = 1.54, *p* =.214, *η_p_*^2^ =.063. These findings suggest that the effect on backward movement was not merely the result of general head motion.

#### 3.2.4. Relationship between head movement and ratings

We examined the within-participant associations between each rating and backward displacement by reference to repeated-measures correlations. Greater backward displacement, as represented by more negative X values, was associated with greater perceived anger (*r_rm_* = -.270, *p* <.001) and greater discomfort (*r_rm_* = -.368, *p* <.001) (Figure 9). Backward displacement was therefore associated with both ratings and provided partial—but not specific—behavioral convergence with subjective discomfort.

**Figure 9.**
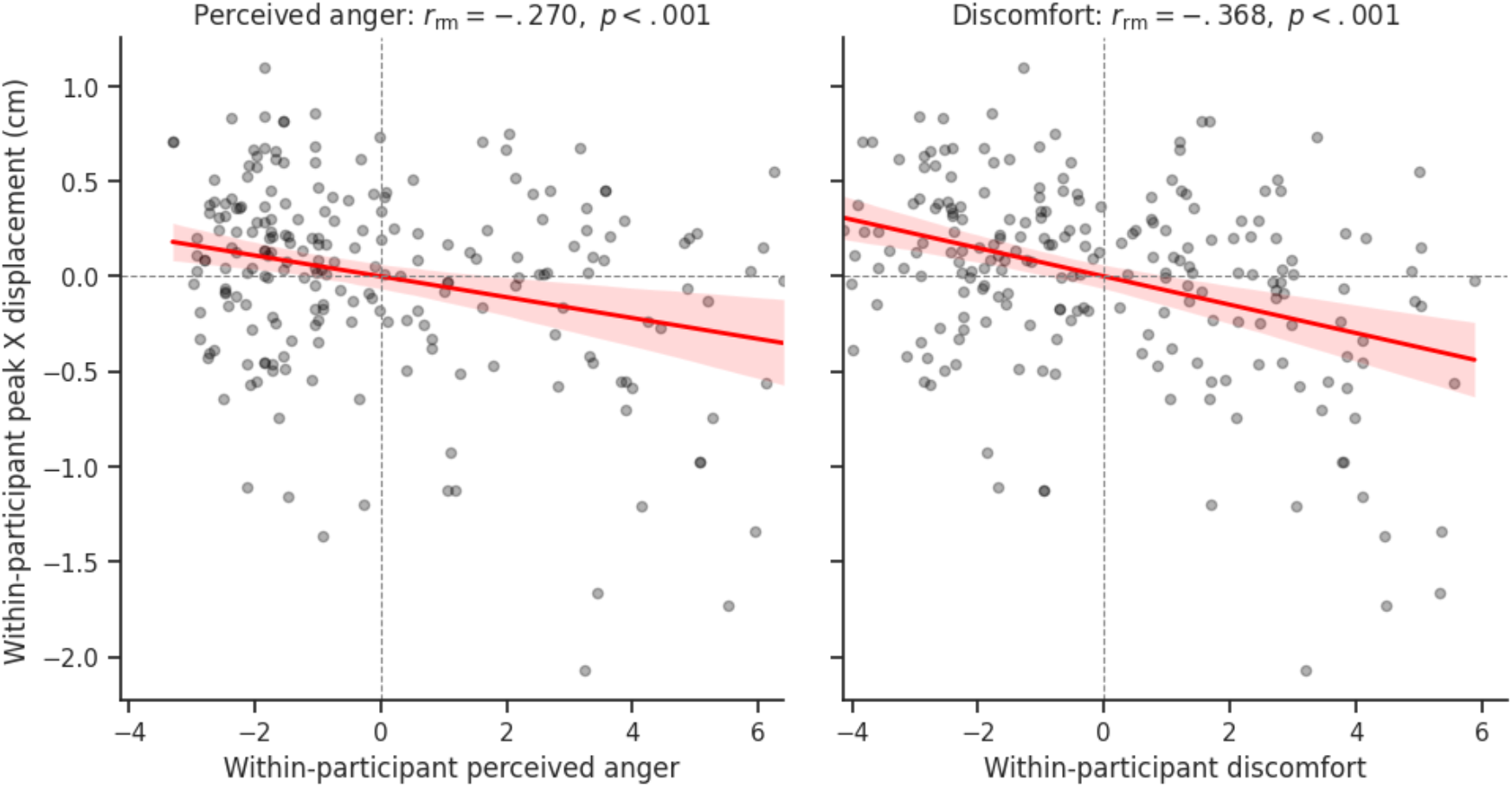
Experiment 2: Repeated-measures associations between backward head displacement and perceived anger and discomfort in Experiment 2. The points represent participant-mean-centered condition-level observations, and the lines indicate linear fits to the centered data. More negative centered X-displacement values indicate greater backward displacement relative to each participant’s mean.

### 3.3. Discussion

In Experiment 2, participants rated the perceived anger and discomfort associated with angry faces, neutral faces, and a featureless mannequin at three distances, and their head movements were recorded. The results reproduced and extended the distinct response patterns that were observed in Experiment 1. Perceived anger was once again driven by face type. Only the angry face was associated with high perceived-anger ratings, whereas the neutral face and the mannequin did not differ. Discomfort was also driven mainly by distance but also exhibited an overall face type effect: angry > neutral > mannequin.

The key comparison in this context was between the neutral face and the mannequin. In comparison with the mannequin, the neutral face elicited greater discomfort at 30 and 100 cm, despite their similarly low perceived-anger ratings. This pattern is consistent with the contribution of a human facial appearance to discomfort beyond the level of perceived anger. However, because the two stimuli also differed in terms of facial features and visual appearance, the contrast should not be interpreted as a direct measure of social or human presence. Head movement exhibited partial behavioral convergence with the ratings. At the closest distance, the backward displacement followed the same order as discomfort—angry face, neutral face, and mannequin—whereas lateral movement did not exhibit any corresponding effects. This direction-specific pattern is broadly consistent with avoidance-related bodily responding (Stins et al., 2011). Finally, backward displacement was associated with both discomfort and perceived anger, thus suggesting that the bodily response was related to the overall evaluation of the encounter rather than specifically to discomfort.

However, in Experiments 1 and 2, the avatar always looked directly at the participant. For an angry face, it was therefore unclear whether the threat was conveyed by the expression itself or by the direct gaze. To examine this point, in Experiment 3, we manipulated gaze direction jointly with expression and distance.

### 4. Experiment 3

Experiment 3 examined the possibility that social discomfort is further shaped by self-relevance, specifically by using gaze direction as an indirect cue. If discomfort is solely a consequence of perceived anger, gaze direction should have negligible effect when the facial expression remains unchanged. Alternatively, if self-relevance contributes to this impact independently, direct gaze should increase discomfort without necessarily increasing perceived anger. In Experiments 1 and 2, the avatar always looked directly at the participant. The question of whether the discomfort elicited by an angry face reflected the facial expression itself or the fact that the observer was the apparent target of the anger remained unanswered. To address this question, Experiment 3 manipulated gaze direction (direct vs. averted) alongside facial expression and interpersonal distance. We hypothesized that gaze direction would have negligible effect on perceived anger because it does not alter the facial expression itself, whereas a direct gaze would increase discomfort by rendering the potential threat more personally relevant.

### 4.1. Methods

#### 4.1.1. Participants

Twenty-four participants were involved in this experiment (21 males and 3 females; all Asian). Mean age = 22.1 years, *SD* = 1.3. Among these individuals, 17 participants had also completed Experiment 2. All other participant details, including the consent procedure and ethics approval, were the same as those described for Experiment 1.

#### 4.1.2. Apparatus

The apparatus used in this experiment was identical to that employed in Experiment 2. As was the case previously, head movements were logged in six degrees of freedom throughout each trial.

#### 4.1.3. Stimuli

The same angry and neutral avatars were used, including the same MetaHuman faces that were employed in Experiments 1 and 2. For each face, the direction of the eyes was manipulated between direct (looking at the participant) and averted (see Figure 10). The averted gaze was presented in two forms, i.e., averted to the left and averted to the right. Only the eyes changed between the direct and averted versions; the face and its position were otherwise identical. As in the previous experiments, each stimulus was presented at eye level at one of three distances (30, 100, or 300 cm).

**Figure 10.**
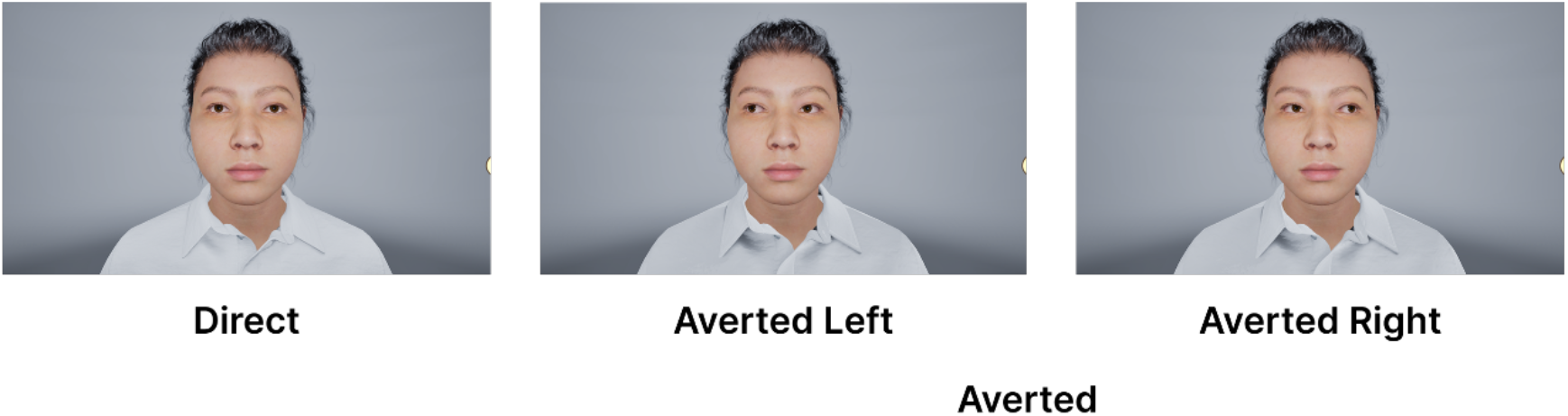
Examples of the direct, averted left, and averted right gaze stimuli used in Experiment 3. Left-averted and right-averted trials were pooled into a single averted condition for analysis.

#### 4.1.4. Procedure

The procedure used in this experiment was identical to that employed for Experiment 1, although gaze replaced the color manipulation. The design crossed expression (angry, neutral) × gaze (direct, averted) × distance (30, 100, 300 cm). Each condition was presented across four avatar identities and repeated twice. For the direct gaze conditions, both repetitions used direct gaze; in contrast, for the averted gaze conditions, one repetition featured a left-averted gaze, whereas the other featured a right-averted gaze. This approach yielded 96 trials in total, which were presented in random order. The left-and right-averted trials were pooled into a single averted gaze condition for analysis. In each trial, the stimulus was shown for 10 s, after which the participants rated the perceived anger of the face and their own discomfort, each of which was scored on a scale ranging from 1 to 10, as in Experiments 1 and 2.

#### 4.1.5. Data analysis

For analysis, the left-averted and right-averted trials were pooled into a single averted level, thus yielding a 2 expression × 2 gaze (direct and averted) × 3 distance design. The same analysis procedure and reporting conventions that were employed in Experiment 1 were used, and expression, gaze, and distance were included as within-participant factors. One trial from participant #20, which was an angry-direct trial at 30 cm, was excluded from the head-movement analysis as a tracking artifact. No participants were excluded from the analysis.

### 4.2. Results

#### 4.2.1. Perceived anger

We performed a three-way repeated-measures ANOVA on the perceived-anger ratings (Figure 11; Table S8). A significant main effect of expression was observed, *F*(1,23) = 583.99, *p* <.001, *η_p_*^2^ =.962. Angry faces were rated as much angrier than neutral faces were, *t*(23) = 24.17, *adj*. *p* <.001, *d* = 4.93. The main effect of distance was also significant, *F*(1.30,29.90) = 89.17, *p* <.001, *η_p_*^2^ =.795, as all three distances differed, such that closer faces were rated as angrier: 30 vs. 100 cm, *d* = 1.32; 30 vs. 300 cm, *d* = 2.12; 100 vs. 300 cm, *d* = 2.31; all *adj*. *p* <.001.

**Figure 11.**
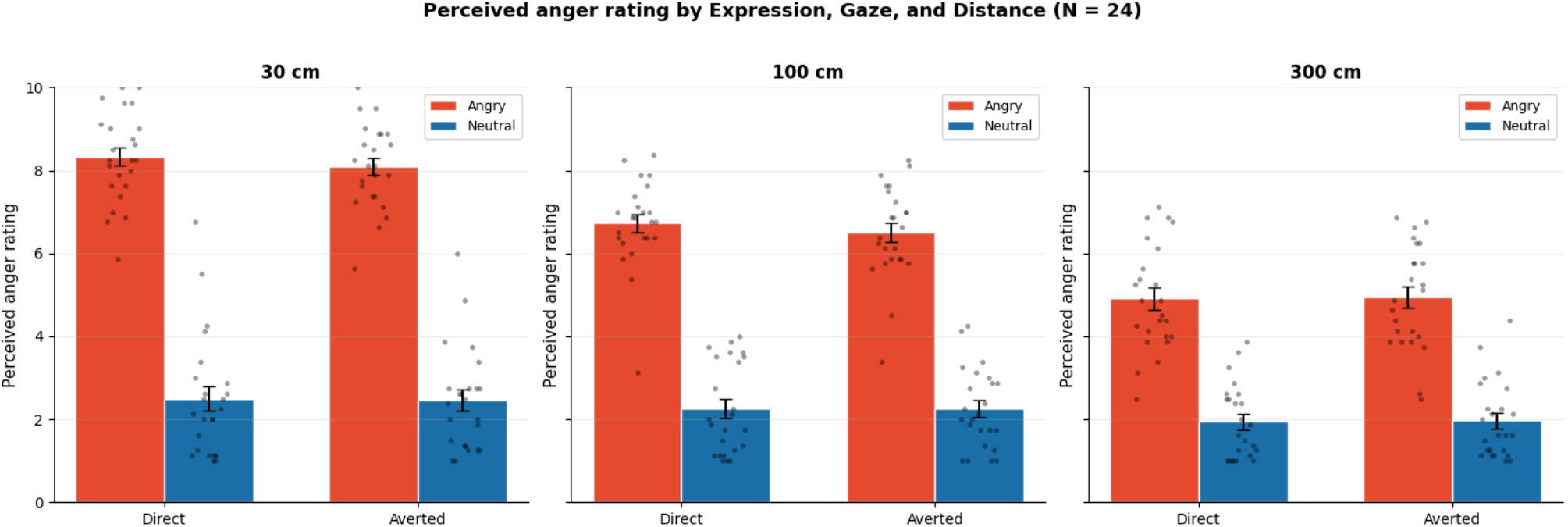
Perceived-anger ratings in Experiment 3 as a function of expression, gaze, and distance. The bars represent condition means, the error bars represent standard errors, and the points represent individual participant values.

Additionally, the expression × distance interaction was significant, *F*(1.42,32.77) = 71.82, *p* <.001, *η_p_*^2^ =.757. In contrast, gaze had no effect on perceived anger, *F*(1,23) = 0.64, *p* =.433, *η_p_*^2^ =.027. Gaze did not interact with expression, *F*(1,23) = 1.62, *p* =.216, *η_p_*^2^ =.066; with distance, *F*(2,46) = 1.27, *p* =.290, *η_p_*^2^ =.052, or with both expression and distance, *F*(2,46) = 0.85, *p* =.434, or *η_p_*^2^ =.036. Accordingly, perceived anger was driven by expression and distance and was unaffected by where the avatar was looking.

#### 4.2.2. Discomfort

The same three-way ANOVA was performed on the discomfort ratings (Figure 12; Table S9). The main effect of distance was the strongest, *F*(1.37,31.59) = 210.91, *p* <.001, *η_p_*^2^ =.902, as closer encounters were rated as more uncomfortable: 30 vs. 100 cm, *t*(23) = 13.31, *d* = 2.72; 30 vs. 300 cm, *t*(23) = 16.03, *d* = 3.27; 100 vs. 300 cm, *t*(23) = 10.80, *d* = 2.21; all *ajd*. *p* <.001.

**Figure 12.**
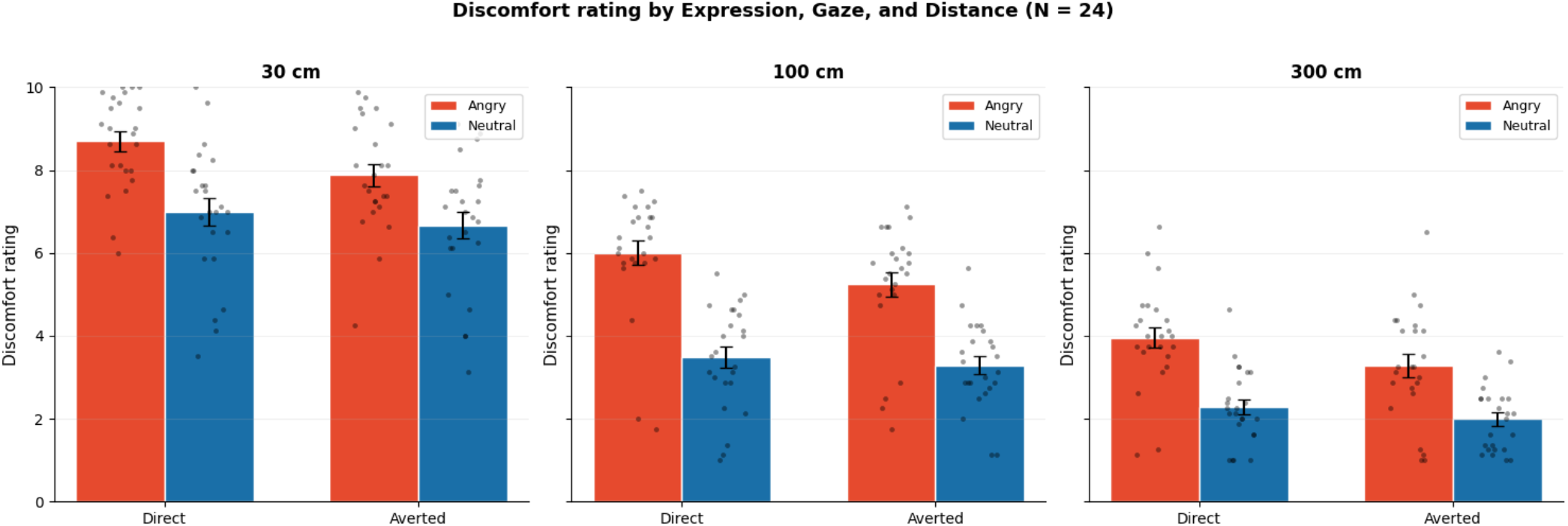
Discomfort ratings in Experiment 3 as a function of expression, gaze, and distance. The bars represent condition means, the error bars represent standard errors, and the points represent individual participant values.

Additionally, the main effect of expression was significant: *F*(1,23) = 90.31, *p* <.001, *η_p_*^2^ =.797. Angry faces were rated as more uncomfortable than neutral faces were, *t*(23) = 9.50, *d* = 1.94. Unlike in the case of perceived anger, the main effect of gaze was significant, *F*(1,23) = 17.13, *p* <.001, *η_p_*^2^ =.427. Direct gaze was rated as more uncomfortable than averted gaze was, *t*(23) = 4.14, *adj*. *p* <.001, *d* = 0.85.

Moreover, the expression × gaze interaction was significant, *F*(1,23) = 25.26, *p* <.001, *η_p_*^2^ =.523. Follow-up comparisons revealed that the direct-versus-averted contrast was significant for angry faces (*t*(23) = 5.67, *adj*. *p* <.001, *d* = 1.16) but did not reach the level of significance for neutral faces (*t*(23) = 2.02, *adj*. *p* =.055, *d* = 0.41). Thus, the effect of gaze on discomfort was stronger for angry faces than for neutral faces. The expression × distance interaction was also significant, *F*(2,46) = 9.32, *p* <.001, *η_p_*^2^ =.288. In contrast, the gaze × distance interaction, *F*(2,46) = 0.17, *p* =.840, *η_p_*^2^ =.008, and the expression × gaze × distance interaction, *F*(2,46) = 0.35, *p* =.706, *η_p_*^2^ =.015, were not significant.

#### 4.2.3. Head movement

Backward and lateral movements were analyzed by conducting a three-way ANOVA. Unlike in Experiment 2, backward movement had no effects. No main effects of expression, *F*(1,23) = 0.14, *p* =.708, *η_p_*^2^ =.006, Gaze, *F*(1,23) = 0.01, *p* =.920, *η_p_*^2^ =.000, or distance, *F*(2,46) = 0.63, *p* =.535, *η_p_*^2^ =.027, were observed. The backward-movement effect observed in Experiment 2 was not observed in Experiment 3. Because the two experiments differed in several respects, this null result does not by itself identify the source of the effect observed in Experiment 2. Additionally, lateral movement did not exhibit any meaningful main effects of expression or gaze and no significant interaction, but there was a significant main effect of distance *F*(1.57,36.17) = 8.27, *p* =.002, *η_p_*^2^ =.264.

### 4.3. Discussion

In Experiment 3, participants rated the perceived anger and discomfort associated with angry and neutral faces with a direct or averted gaze at three distances, and their head movement was recorded. The results indicated that gaze affected feeling but not perception. Perceived anger was driven by expression and distance and was unaffected by gaze, whereas discomfort was increased by direct gaze, specifically for the angry face. These findings extend the distinct response patterns observed in the previous experiments. Gaze, like distance in Experiment 1 and the human–face–mannequin contrast in Experiment 2, changed how uncomfortable the encounter felt without changing how the face was perceived.

The expression × gaze interaction represented the main result. Direct gaze increased discomfort more strongly for angry faces than for neutral faces. This pattern is consistent with the interpretation that the direct gaze increased the self-relevance of the angry expression, although self-relevance was not measured directly. This interpretation is consistent with accounts in which direct gaze increases self-relevance and signals that a threat is directed at the observer (Adams & Kleck, 2005; Hietanen et al., 2008; Ewbank et al., 2009). The absence of any gaze effect on perceived anger is notable because some previous studies have reported that gaze direction can shift perceptions of facial emotions, in which context direct gaze enhances perceptions of anger (Adams & Kleck, 2005; Ewbank et al., 2009; McCrackin & Itier, 2019). In the present data, the angry face was assigned similar perceived-anger ratings whether it looked at the participant or away. We suggest that, in these conditions, gaze acted on the affective response rather than on the perceptual judgment.

Unlike in Experiment 2, neither backward nor lateral head movement had reliable effects in Experiment 3, a part from a significant main effect of distance for lateral movement. This difference may reflect the absence of the human–face–mannequin contrast, but the present data do not allow us to determine why the head-movement effect was observed in Experiment 2 but not in Experiment 3. Head movement therefore exhibited only partial and experiment-specific convergence with the subjective ratings.

Taken together, these three experiments reveal that perception and feeling can be separated by distance, facial stimulus type, and gaze.

## 5. General Discussion

### 5.1. Summary of Findings

We conducted three experiments to test a single question concerning whether perceived anger and subjective discomfort in a social encounter are driven by the same cues or whether they can be separated. In Experiment 1, we tested whether perceived anger is driven mainly by expression and discomfort is driven mainly by distance. In Experiment 2, we examined whether discomfort differed between human facial stimuli and a featureless mannequin. In Experiment 3, we investigated whether gaze direction modulates feeling without changing perception. Across these three experiments, perceived anger was driven mainly by expression and increased with both increasing distance and red color. Discomfort, in contrast, was driven mainly by distance, was increased by expression and red color, differed between the human-face and mannequin conditions, and was modulated by direct gaze. These findings indicate that what a face is perceived to be and how uncomfortable it feels are separable: a neutral human face increased discomfort relative to a mannequin without increasing perceived anger, a direct gaze increased discomfort without changing perceived anger, and these two ratings were dominated by varied factors, i.e., expression for perceived anger and distance for discomfort.

### 5.2. Perceived Anger and Discomfort Are Separable

The results demonstrate distinct patterns of sensitivity between perceived anger and discomfort judgments. Perceived anger was dominated by the face itself: angry faces were rated higher in terms of perceived anger than neutral faces were in every experiment, and red color increased perceived anger in Experiment 1. However, two cues that raised discomfort left perceived anger largely unchanged: a neutral face and a featureless mannequin were rated similarly low in terms of perceived anger in Experiment 2, and an angry face was assigned similar perceived-anger ratings whether it looked at the participant or away in Experiment 3. Discomfort differed between the human-face and mannequin conditions and was increased by a direct gaze. We suggest that perceived anger reflects a judgment concerning the facial signal, whereas discomfort reflects the encounter as a whole. This interpretation is in line with the proposal that close interpersonal distance elicits discomfort and arousal through mechanisms that do not require a threatening face (Hayduk, 1983; Kennedy et al., 2009; Candini et al., 2021). This research also extends that proposal by demonstrating that discomfort also differed as a function of facial stimulus type and gaze direction.

### 5.3. Why Distance, Facial Stimulus Type, and Gaze Matter

This pattern may be due to the fact that the cues do not carry the same kind of information. Expression and facial color mainly specify what the face is expressing. Distance, facial stimulus type, and gaze provide information about different aspects of the encounter. Recent studies have also suggested that emotion perception can vary with the spatial position of a face and with approach-avoidance behavior (Tamura et al., 2026; Kobayashi et al., 2025). A close neutral face may be uncomfortable because it enters one’s personal space (Hayduk, 1983; Candini et al., 2021), even if it does not signal anger. A direct angry gaze may be uncomfortable because the threat appears to be directed at the observer (Adams & Kleck, 2005; Hietanen et al., 2008), even if the expression itself is not judged as angrier. In this sense, discomfort may depend less on the face alone and more on the relationships among the face, the body, and the observer. This interpretation is consistent with previous research on interpersonal distance and intimacy regulation (Hall, 1966; Argyle & Dean, 1965) and with studies that have demonstrated that gaze direction changes the social meaning of facial expressions (Ewbank et al., 2009; McCrackin & Itier, 2019).

### 5.4. Head Movement as a Bodily Index

Head movement served as an additional behavioral index of the encounter, although its effects were less consistent than those that were observed in the ratings. In Experiment 2, participants exhibited greater backward displacement in response to human faces than the mannequin, particularly at the closest distance. This pattern is broadly consistent with previous evidence indicating that social and emotional cues can elicit avoidance-related bodily responses, including backward movement away from angry faces (Stins et al., 2011; Welsch et al., 2023). More broadly, social threat can modulate bodily behavior in several ways. For example, angry faces have been reported to reduce body sway and elicit freeze-like responses (Roelofs et al., 2010), and freezing has been conceptualized as an active preparatory state that supports subsequent threat-coping behavior rather than as simple passive immobility (Roelofs & Dayan, 2022). The backward displacement observed in this context should not be interpreted as a freezing response, but it provides further evidence indicating that social cues can influence bodily regulation during an encounter. However, this effect was not replicated in Experiment 3, in which all the stimuli were human faces, and backward displacement in Experiment 2 was associated with both perceived anger and discomfort. Head movement should therefore be viewed as a form of partial behavioral convergence rather than as a specific measure of social discomfort.

### 5.5. Limitations and Directions for Future Research

This study has several limitations. First, the avatars appeared realistic and had full bodies, but they did not move during the trial. A moving avatar that approaches or moves away may elicit stronger or different responses (Bailenson et al., 2003; Bönsch et al., 2018; Ruggiero et al., 2017). Second, all the participants in this research were East Asian adults. Since interpersonal distance differs across cultures, the present results may not be the same for other groups (Hall, 1966; Sorokowska et al., 2017). Third, head movement is only a simple bodily measure, and the ratings were provided in response to single questions. Future studies should include other measures, such as physiological arousal, to support this interpretation (Candini et al., 2021).

Future research can build on these findings in several ways. The clearest of these ways follows from the central result of this research: if perception and feeling are separable, the size of the separation may differ among individuals. A further direction involves examining individual differences in autistic traits. Previous studies have reported differences in interpersonal distance regulation in cases involving autism (Asada et al., 2016; Gessaroli et al., 2013). Future studies could test whether autistic traits are related to the separation between perceived anger and discomfort in virtual social encounters. A second direction involves movement: in all three experiments, the avatar was static, and a moving avatar that approaches or withdraws would allow the dynamics of such responses to be measured. A third direction focuses on the inclusion of additional control stimuli matched to the human faces in terms of shape, texture, and visual complexity. Such controls could help distinguish the contribution of human facial appearance from lower-level visual differences between the faces and the mannequin.

## 6. Conclusion

We investigated whether the perception of a face and the discomfort that it produces are driven by the same cues across three virtual reality experiments. Perceived anger was driven mainly by expression; in contrast, discomfort was driven mainly by distance, differed between the human-face and mannequin conditions, and was modulated by gaze direction. Two dissociations were particularly informative: in comparison with the mannequin, the neutral human faces elicited greater discomfort despite similarly low perceived-anger ratings, and direct gaze increased discomfort without changing perceived anger. Head movement exhibited partial behavioral convergence with greater backward displacement from human faces than from the mannequin. Together, these findings suggest that a social encounter should be treated not as a single response but rather as a combination of perceptual, affective, and bodily components that follow different cues.

## Acknowledgments

This work was supported by JSPS KAKENHI (Grant Numbers JP25K21323 to H.T., JP23KK0183 to T.M. and JP25H01141 to S.N.), JST Grant Number JPMJPF2502, and the Foundation of Amano Institute of Technology.

## Data and code availability

The data and code that support the findings of this study are openly available at the Open Science Framework repository (https://osf.io/fqczw).

## Declaration of the use of Generative AI and AI-assisted technologies in the writing process

During the preparation of this work, the authors used ChatGPT 5.6 Sol to improve the language of the manuscript, which has also been proofread by native English speakers through an English editing service. After the use of this tool and service, the authors reviewed and edited the content as needed and take full responsibility for the content of the publication.

