## Supplementary Information for "What makes an angry face uncomfortable? Distinct contributions of facial expression, interpersonal distance, facial stimulus type, and gaze in virtual reality"

Table S1. Experiment 1. Results of the three-way repeated-measures ANOVA for perceived-anger ratings.

| Effect | *F* | *df* | *p* | $\eta p^2$ |
|---|---|---|---|---|
| Expression | 582.07 | 1, 23 | < .001 | .962 |
| Color | 29.95 | 1, 23 | < .001 | .566 |
| Distance | 73.81 | 1.21, 27.86 | < .001 | .762 |
| Expression × Color | 0.37 | 1, 23 | .547 | .016 |
| Expression × Distance | 49.37 | 2, 46 | < .001 | .682 |
| Color × Distance | 1.89 | 2, 46 | .163 | .076 |
| Expression × Color × Distance | 3.27 | 2, 46 | .047 | .125 |

Table S2. Experiment 1. Post hoc tests for the effect of the three-way interaction on perceived anger

| Condition | Comparison | *t(23)* | *adj. p* | *d* |
|---|---|---|---|---|

| | | | | |
|---|---|---|---|---|
| Angry, 30 cm | Red > Natural | 3.74 | .006 | 0.76 |
| Angry, 100 cm | Red > Natural | 3.25 | .021 | 0.66 |
| Angry, 300 cm | Red > Natural | 6.17 | < .001 | 1.26 |
| Neutral, 30 cm | Red > Natural | 3.76 | .006 | 0.77 |
| Neutral, 100 cm | Red > Natural | 1.64 | .688 | 0.33 |
| Neutral, 300 cm | Red > Natural | 3.36 | .016 | 0.69 |

Table S3. Experiment 1. Results of the three-way repeated-measures ANOVA for discomfort ratings.

| Effect | *F* | *df* | *p* | $\eta p^2$ |
|---|---|---|---|---|
| Expression | 79.96 | 1, 23 | < .001 | .777 |
| Color | 18.05 | 1, 23 | < .001 | .440 |
| Distance | 236.47 | 2, 46 | < .001 | .911 |
| Expression × Color | 4.33 | 1, 23 | .049 | .159 |
| Expression × Distance | 4.25 | 2, 46 | .020 | .156 |
| Color × Distance | 2.75 | 2, 46 | .075 | .107 |
| Expression × Color × Distance | 3.08 | 2, 46 | .055 | .118 |

Table S4. Results of the two-way repeated-measures ANOVA for perceived-anger ratings.

| Effect | *F* | *df* | *p* | $\eta p^2$ |
|---|---|---|---|---|
| Face Type | 144.04 | 1.52, 34.86 | < .001 | .862 |
| Distance | 64.90 | 1.16, 26.59 | < .001 | .738 |
| Face Type × Distance | 27.26 | 1.88, 43.28 | < .001 | .542 |

Table S5. Results of the two-way repeated-measures ANOVA (face type × distance) on discomfort ratings.

| Effect | *F* | *df* | *p* | $\eta p^2$ |
|---|---|---|---|---|
| Face Type | 75.11 | 1.47, 33.73 | < .001 | .766 |
| Distance | 147.59 | 1.51, 34.77 | < .001 | .865 |
| Face Type × Distance | 5.75 | 1.66, 38.09 | .010 | .200 |

Table S6. Results of the two-way repeated-measures ANOVA for peak head movement (backward and lateral movements)

| Axis | *Effect* | *F* | *df* | *p* | $\eta p^2$ |
|---|---|---|---|---|---|
| Backward (X) | Face Type | 16.51 | 2, 46 | < .001 | .418 |
| | Distance | 5.33 | 1.39,32.04 | .018 | .188 |
| | Face Type × Distance | 4.68 | 2.79,64.21 | .006 | .169 |

Table S7. Post hoc comparisons for discomfort and backward head movement in Experiment 2.

| Comparison | *t(23)* | *adj. p* | *d* |
|---|---|---|---|
| Discomfort by Face Type | | | |
| Angry vs. Neutral | 9.42 | < .001 | 1.92 |
| Angry vs. Mannequin | 9.41 | < .001 | 1.92 |
| Neutral vs. Mannequin | 4.13 | .001 | 0.84 |
| Backward Movement (X) by Face Type | | | |
| Angry vs. Neutral | -2.29 | .095 | -0.47 |
| Angry vs. Mannequin | -5.33 | < .001 | -1.09 |
| Mannequin vs. Neutral | 3.73 | .003 | 0.76 |
| Backward Movement (X) at 30 cm | | | |
| Angry vs. Neutral | -3.23 | .011 | -0.66 |
| Angry vs. Mannequin | -4.76 | < .001 | -0.97 |
| Mannequin vs. Neutral | 3.53 | .005 | 0.72 |

Table S8. Results of the three-way repeated-measures ANOVA for perceived-anger ratings in Experiment 3

| Effect | *F* | *df* | *p* | $\eta p^2$ |
|---|---|---|---|---|
| Expression | 583.99 | 1, 23 | < .001 | .962 |
| Gaze | 0.64 | 1, 23 | .433 | .027 |

| | | | | |
|---|---|---|---|---|
| Distance | 89.17 | 1.30, 29.90 | < .001 | .795 |
| Expression × Gaze | 1.62 | 1, 23 | .216 | .066 |
| Expression × Distance | 71.82 | 1.42, 32.77 | < .001 | .757 |
| Gaze × Distance | 1.27 | 2, 46 | .290 | .052 |
| Expression × Gaze × Distance | 0.85 | 2, 46 | .434 | .036 |

Table S9. Results of the three-way repeated-measures ANOVA for discomfort ratings in Experiment 3

| Effect | *F* | *df* | *p* | $\eta p^2$ |
|---|---|---|---|---|
| Expression | 90.31 | 1, 23 | < .001 | .797 |
| Gaze | 17.13 | 1, 23 | < .001 | .427 |
| Distance | 210.91 | 1.37, 31.59 | < .001 | .902 |
| Expression × Gaze | 25.26 | 1, 23 | < .001 | .523 |
| Expression × Distance | 9.32 | 2, 46 | < .001 | .288 |
| Gaze × Distance | 0.17 | 2, 46 | .840 | .008 |
| Expression × Gaze × Distance | 0.35 | 2, 46 | .706 | .015 |
